# On spatial molecular arrangements of SARS-CoV2 genomes of Indian patients

**DOI:** 10.1101/2020.05.01.071985

**Authors:** Sk. Sarif Hassan, Atanu Moitra, Ranjeet Kumar Rout, Pabitra Pal Choudhury, Prasanta Pramanik, Siddhartha Sankar Jana

## Abstract

A pandemic caused by the SARS-CoV2 is being experienced by the whole world since December, 2019. A thorough understanding beyond just sequential similarities among the protein coding genes of SARS-CoV2 is important in order to differentiate or relate to the other known CoVs of the same genus. In this study, we compare three genomes namely MT012098 (India-Kerala), MT050493 (India-Kerala), MT358637 (India-Gujrat) from India with NC_045512 (China-Wuhan) to view the spatial as well as molecular arrangements of nucleotide bases of all the genes embedded in these four genomes. Based on different features extracted for each gene embedded in these genomes, corresponding phylogenetic relationships have been built up. Differences in phylogenetic tree arrangement with individual gene suggest that three genomes of Indian origin have come from three different origins or the evolution of viral genome is very fast process. This study would also help to understand the virulence factors, disease pathogenicity, origin and transmission of the SARS-CoV2.

## 1. Introduction

The disease COVID-19 is caused by the SARS-CoV2 initiated in late December 2019 in Wuhan, China, and since then it has been impulsed various countries across the world [1]. Presently, this disease, a pandemic as announced by the WHO, is a major health concern [2]. The family of coronaviruses is enclosed by different CoVs which are a single-stranded, positive-sense RNA genome of size approximately 26-32 kb [3]. The CoVs are classified into four genera, the *α*-CoVs, *β*-CoVs, *γ*-CoVs and *δ*-CoVs [4]. One of most important genera of coronaviruses is the *β*-CoVs where the present SARS-CoV2 belongs [5]. The *β*-CoVs mainly infect humans, bats including other animals such as camels, and rabbits and so on [6]. Two-third of the SARS-CoV2 genomes from 5’ end is conserved for the ORF1 gene which encodes sixteen polyproteins and the 3’ ends contains various structural protein coding genes including surface (S), envelope (E), membrane (M), and nucleocapsid (N) proteins [7]. In addition there are six accessory protein coding genes such as ORF3a, ORF6, ORF7a, ORF7b, and ORF8 also present in the SARS-CoV2 genome [8]. The spatial arrangement of genes over the SARS-CoV2 genome is presented in the Fig.1. [9].

**Figure 1:**
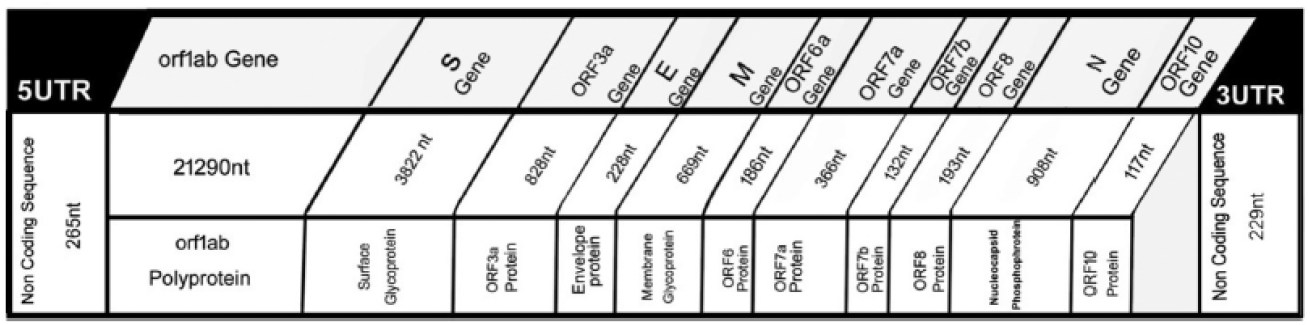
Spatial arrangement of genes over a typical SARS-CoV2 genome, Credit: [9].

The polyprotein ORF1ab encoded by the ORF1 gene play key roles in virus pathogenesis, cellular signalling, modification of cellular gene expression[10]. The envelope (E) proteins play multiple roles during infection, including virus morphogenesis [11, 12]. The N protein encoded by the gene N plays a vital role in the virus morphogenesis and assembly [13, 14]. The M gene encodes the M protein which plays a central role in virus morphogenesis and assembly via its interactions with other viral proteins [15, 16]. It does determine the shapes of the virions, promotes membrane curvature. The M gene sequence of SARS-CoV2 is similar to that of SARS-CoV and MERS-CoV with 90.1% and 39.2% respectively [17]. The S protein (S gene) is one of the most important structural proteins, which is used as a key that the virus uses to enter host cells. The spike protein attaches the virion to the cell membrane by interacting with host receptor and infects the host cell [18, 19]. In viral replication, the accessory proteins such as ORF3a, ORF6, ORF7a, ORF7b, ORF8 have a key role [20].

SARS-COV2 genome shows 79.6% identity with SRS-COV1 genome [21]. It is reported that the Spike glycoprotein of the Wuhan coronavirus is modified via homologous recombination [22]. The SARS-CoV2 is more phylogenetically related to SARS-CoV than to MERS-CoV [23]. Still, the proximal origin of COVID-19 transmission or evolutionary relationship of SARS-CoV2 and other coronaviruses is very much controversial. The outbreak and infectious behaviour of the SARS-CoVs and the lack of effective treatments for CoV infections demand the need of detailed understanding of coronaviral molecular biology, with a specific focus on both their structural proteins as well as their accessory proteins.

From a molecular biology perspective, figuring out why the virus is so much virulent and infectious than other CoVs belonging to the genus *β*-CoVs is one of the most important aspects to look into [24]. The present SARS-CoV2 genomes are classified, based on SNPs, into two broad groups known as L and S [25]. The nature of virulence is also associated with the L type of CoV2 genomes.

Clearly, on having information of sequential similarity among genes and genomes of various CoVs is not enough to decipher the deep message regarding various characteristics viz. virulence, infection and transmission capacities, embedded in the RNA sequence. So in order to find out the genomic information of the two types (L and S) of SARS-CoV2, an attempt is made to discover the molecular and spatial organizations of each gene embedded over a sample of four genomes of which three of them are from India and one from China-Wuhan.

### 1.1. Datasets and Methods

In the NCBI virus database, as on 30th April, 2020, there are three complete genomes viz. MT012098 (India-Kerala), MT050493 (India-Kerala) and MT358637 (India-Gujrat) of SARS-CoV2 from Indian patients are available, which we consider for this present study. As a reference genome, NC_045512 (China-Wuhan) is taken. Note that, the genomes MT012098, MT050493 and NC_045512 belong to the S-type and other genome MT358637 from India belongs to L-type as per classification made based on SNPs[25]. An information regarding the lengths and names (followed strictly by NCBI database) of all eleven genes embedded in the four genomes is presented in the Table 1. Note that, in the Table 1, ‘*’ denotes absence of the gene in the respective genome.

**Table 1:** Information of the eleven genes of the India’s and Wuhan’s Genomes

| Genes | NC_045512 (China) | MT012098 (India) | MT050493 (India) | MT358637 (India) |
| --- | --- | --- | --- | --- |
| ORF1 | 21290 | 21290 | 21290 | 21290 |
| S (ORF2) | 3822 | 3819 | 3822 | 3822 |
| ORF3a | 828 | 828 | 828 | 828 |
| E (ORF4) | 228 | 228 | 228 | 228 |
| M (ORF5) | 669 | 669 | 669 | 669 |
| ORF6 | 186 | 186 | 186 | 186 |
| ORF7a | 366 | 366 | 366 | 366 |
| ORF7b | 132 | * | * | 132 |
| ORF8 | 366 | 366 | 366 | 366 |
| N (ORF9) | 1260 | 1260 | 1260 | 1260 |
| ORF10 | 117 | 117 | 117 | 117 |

It is noted that the gene ORF7b is absent in the genomes MT012098 and MT050493 as shown in Table 1 with ‘*’ mark.

From the sequence based similarity, a phylogenetic relations is given in the Fig.2 which describes that the genomes NC_045512 from Wuhan and MT012098 from India are very close to each other with 99.98% sequential similarity as mentioned in the article by Yadav P.D. et.al. [26].

**Figure 2:**
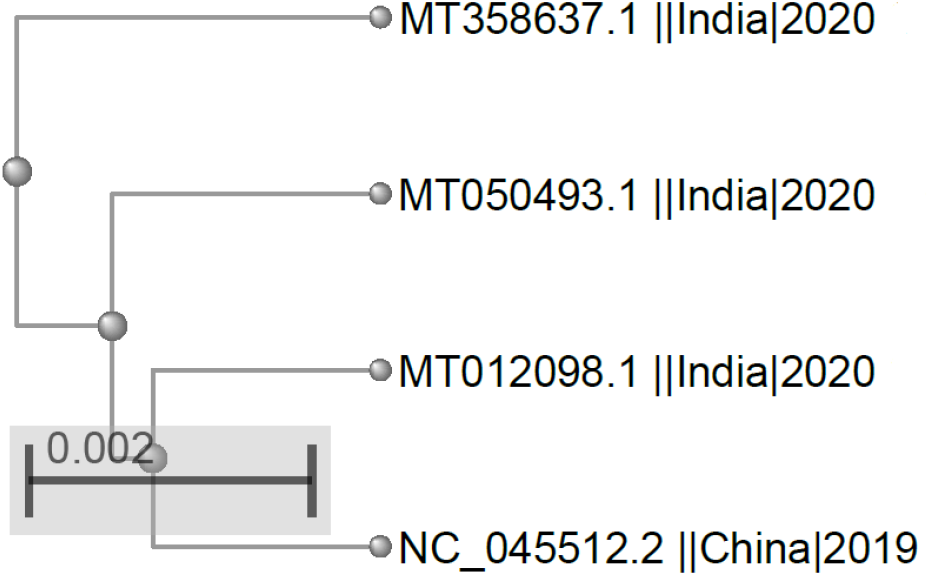
Phylogeny among the four genomes based on sequential similarities. Credit: NCBI

The phylogeny shows that the two genomes namely NC_045512 (China-Wuhan) and MT012098 (India-Kerala) are sequentially similar to the genome MT050493 (India-Kerala) while the genome MT358637 (India-Gujrat) belongs to the other branch of the tree.

At first, each sequence to a binary sequence of 1’*s* and 0’*s* as per the definition 1 has been transformed to a binary representation. Here purine (A,G) and pyrimidine (T,C) bases are represented as ‘1’ and ‘0’ respectively. This binary representation is named as purine-pyrimidine representation [27, 28, 29, 30, 31].

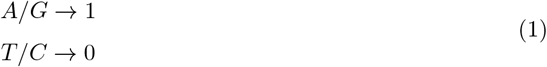

Also each sequence is transformed to a binary sequence with respect to a nucleotide base *B* of 1’*s* and 0’*s* as per the following definition 2.

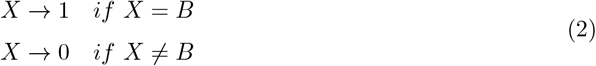

Hence four binary representations for each nucleotide *B* ∈ {*A, T, C, G*} would be derived for a given nucleotide sequence. These binary representations are actually the spatial templates of all the nucleotides. Each of these spatial templates is to be analysed using various methods as mentioned in the following.

#### Binary Shannon Entropy

The Shannon entropy (SE) measures information entropy of a Bernoulli process with probability *p* of the two outcomes (0*/*1) [32, 33]. It is defined as

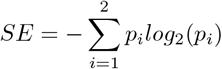

where 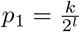 and 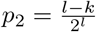; here *l* is the length of the binary sequence and *k* is the number of 1’s in the binary sequence of length *l*. The binary Shannon entropy is a measure of the uncertainty in a binary sequence. If the probability *p* = 0, the event is certainly never to occur, and so there is no uncertainty, leading to an entropy of 0. Similarly, if the probability *p* =1, the result is certain, so the entropy must be 0. When *p* = 0.5, the uncertainty is at a maximum and consequently the SE is 1.

#### Nucleotide Conservation Shannon Entropy

Shannon entropy is a measure of the amount of information (measure of uncertainty). Conservation of each of the four nucleotides has been determined using Shannon entropy [34]. For a given RNA sequence, the conservation SE is calculated as follows: where *p_Ni_* represents the occurrence frequency of a nucleotide *N_i_* in a RNA sequence.

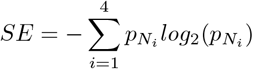

#### Hurst Exponent

The Hurst Exponent (HE) is used to interpret the trend of a sequence, which could be positive or negative [35]. The HE belongs to the unit interval (0, 1). If HE lies within (0, 0.5) then the sequence possesses a negatively trending. Otherwise if the HE belongs to (0.5, 1) then the sequence is positively trending. If the HE is turned out to be 0.5 then the sequence must possesses randomness.

The *HE* of a sequence *b_n_* (length: *n*) is defined as

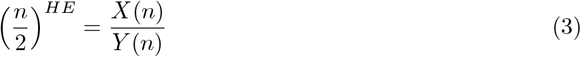

where

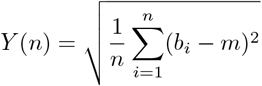

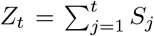 for *t* = 1, 2,3,…, *n* where *S_t_* = *b_t_* – *m* for 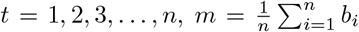 and *X*(*n*) = *max*(*Z_i_*) – *min*(*Z_i_*) for *i* = 1, 2, 3,…, *n*

In addition to these two parameters Shannon entropy and Hurst exponent, some basic derivative features such as nucleotide frequency (Freq), double nucleotide frequency, codon usage frequency, GC content, purine-pyrimidine density are obtained for a given nucleotide sequence [29, 31]. Also based on nucleotide densities, a decreasing order (density order) is obtained for a given sequence. It is worth noting that the first positive frame has been considered to determine codons and double nucleotides over a given gene.

## 2. Results

Important features for the all genes ORF1-10 of the four genome using above methods have been analyzed and interpreted in the following sections.

### 2.1. Quantifications of genes over the four SARS-CoV2 genomes

For a given gene, we define a feature vector as (length, frequency of individual nucleotides, GC content, % of purines and pyrimidines, frequency of each codon usage, frequencies of each double nucleotides, Shannon entropy (SE) and Hurst Exponent (HE) of the spatial representations of each nucleotides and purine-pyrimidine).

Here we briefly state the findings based on the feature vectors derived for every gene embedded in the four genomes and accordingly based on the findings some discussions are made. Before we proceed to make specific observations about codon and double nucleotide usages we present a table (Table 2) below describing the molecular information of the each genes with associated remarks.

**Table 2:**
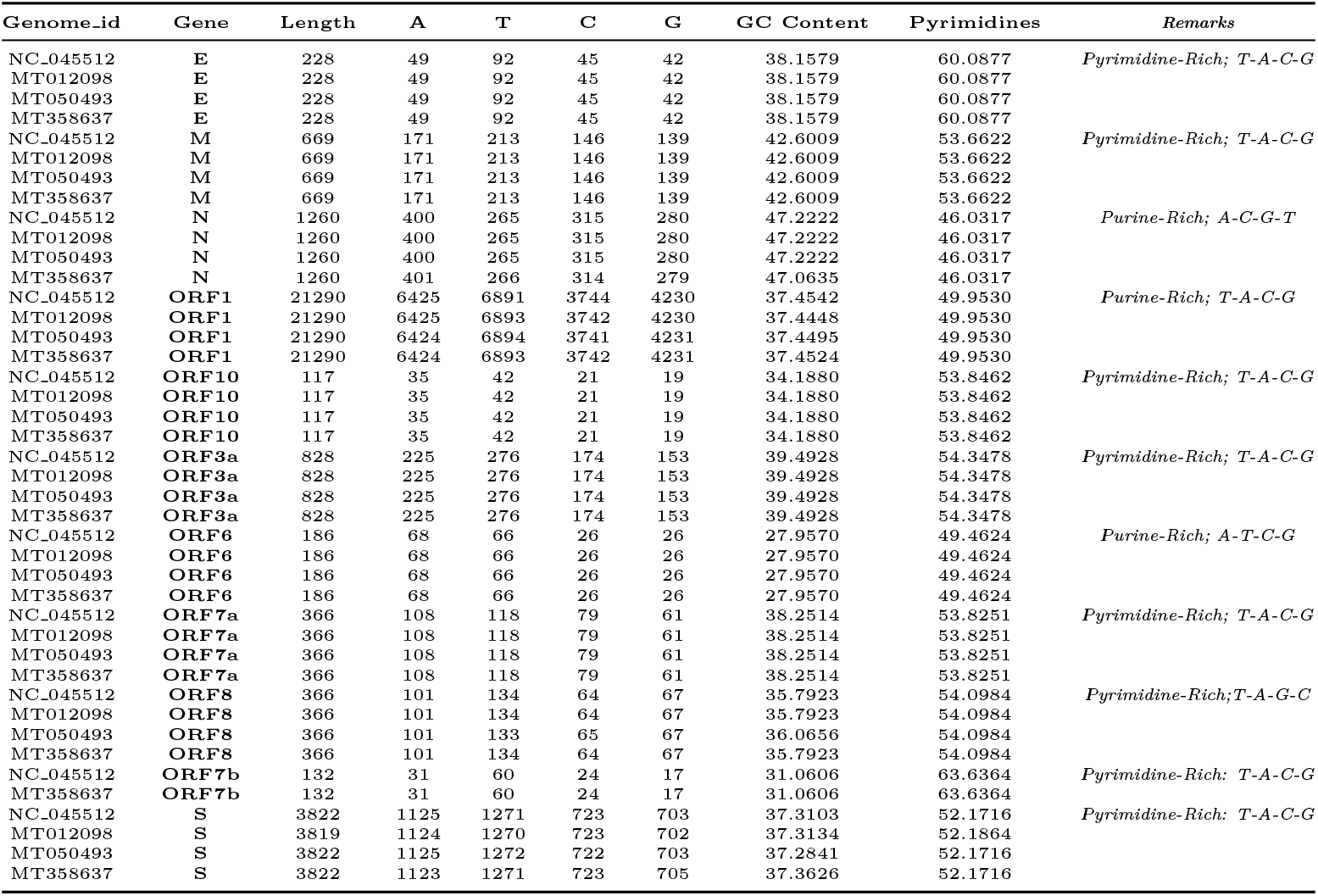
The molecular descriptions of the genes of four genomes with remarks

In Table 3 below, SEs and HEs of spatial representations of the four nucleotide bases as well as of the purine-pyrimidine representations over each genes of four different genomes, are presented. Here HEN and SEN denote the Hurst exponent and Shannon entropy respectively of the spatial representation of the nucleotide N. The B_HE and BSE represent the Hurst exponent and Shannon entropy of the spatial representation of the purine-pyrimidine bases. SE_Conv stands for the Shannon entropy of the conservation of nucleotides over a gene.

**Table 3:** The SEs and HEs of the spatial representations of the four nucleotide bases as well as of the purine-pyrimidine representations over the genes of four different genomes

| Genome_id | Gene | HE_A | HE_C | HE_T | HE_G | SE_A | SE_C | SE_T | SE_G | B_HE | B_SE | SE-Conv |
| --- | --- | --- | --- | --- | --- | --- | --- | --- | --- | --- | --- | --- |
| NC_045512 | E | 0.63928 | 0.54726 | 0.55748 | 0.57277 | 0.75077 | 0.71663 | 0.97297 | 0.68920 | 0.62451 | 0.97044 | 0.97673 |
| MT012098 | E | 0.63928 | 0.54726 | 0.55748 | 0.57277 | 0.75077 | 0.71663 | 0.97297 | 0.68920 | 0.62451 | 0.97044 | 0.97903 |
| MT050493 | E | 0.63928 | 0.54726 | 0.55748 | 0.57277 | 0.75077 | 0.71663 | 0.97297 | 0.68920 | 0.62451 | 0.97044 | 0.97903 |
| MT358637 | E | 0.63928 | 0.54726 | 0.55748 | 0.57277 | 0.75077 | 0.71663 | 0.97297 | 0.68920 | 0.62451 | 0.97044 | 0.98152 |
| NC_045512 | M | 0.63822 | 0.62440 | 0.66460 | 0.57028 | 0.82004 | 0.75693 | 0.90262 | 0.73720 | 0.63096 | 0.99613 | 0.97735 |
| MT012098 | M | 0.63822 | 0.62440 | 0.66460 | 0.57028 | 0.82004 | 0.75693 | 0.90262 | 0.73720 | 0.63096 | 0.99613 | 0.97903 |
| MT050493 | M | 0.63822 | 0.62440 | 0.66460 | 0.57028 | 0.82004 | 0.75693 | 0.90262 | 0.73720 | 0.63096 | 0.99613 | 0.97903 |
| MT358637 | M | 0.63822 | 0.62440 | 0.66460 | 0.57028 | 0.82004 | 0.75693 | 0.90262 | 0.73720 | 0.63096 | 0.99613 | 0.98152 |
| NC_045512 | N | 0.61742 | 0.50535 | 0.53219 | 0.55515 | 0.90160 | 0.81128 | 0.74209 | 0.76420 | 0.57043 | 0.99545 | 0.97929 |
| MT012098 | N | 0.61742 | 0.50535 | 0.53219 | 0.55515 | 0.90160 | 0.81128 | 0.74209 | 0.76420 | 0.57043 | 0.99545 | 0.97903 |
| MT050493 | N | 0.61742 | 0.50535 | 0.53219 | 0.55515 | 0.90160 | 0.81128 | 0.74209 | 0.76420 | 0.57043 | 0.99545 | 0.97903 |
| MT358637 | N | 0.61484 | 0.51273 | 0.53645 | 0.55913 | 0.90247 | 0.81002 | 0.74360 | 0.76277 | 0.57043 | 0.99545 | 0.98152 |
| NC_045512 | S | 0.59665 | 0.59186 | 0.61746 | 0.55111 | 0.87427 | 0.69973 | 0.91751 | 0.68861 | 0.58340 | 0.99864 | 0.97644 |
| MT012098 | S | 0.59687 | 0.59037 | 0.61752 | 0.55202 | 0.87423 | 0.70004 | 0.91751 | 0.68836 | 0.58340 | 0.99862 | 0.97903 |
| MT050493 | S | 0.59665 | 0.59070 | 0.61623 | 0.55111 | 0.87427 | 0.69918 | 0.91777 | 0.68861 | 0.58340 | 0.99864 | 0.97903 |
| MT358637 | S | 0.59926 | 0.59186 | 0.61746 | 0.54722 | 0.87361 | 0.69973 | 0.91751 | 0.68973 | 0.58340 | 0.99864 | 0.98152 |
| NC_045512 | ORF1 | 0.62093 | 0.56846 | 0.66817 | 0.59449 | 0.88346 | 0.67093 | 0.90833 | 0.71929 | 0.64206 | 1.00000 | 0.97627 |
| MT012098 | ORF1 | 0.62093 | 0.56652 | 0.66840 | 0.59449 | 0.88346 | 0.67072 | 0.90843 | 0.71929 | 0.64206 | 1.00000 | 0.97903 |
| MT050493 | ORF1 | 0.62075 | 0.56849 | 0.66828 | 0.59536 | 0.88340 | 0.67062 | 0.90848 | 0.71938 | 0.64206 | 1.00000 | 0.97903 |
| MT358637 | ORF1 | 0.62075 | 0.56800 | 0.66793 | 0.59427 | 0.88340 | 0.67072 | 0.90843 | 0.71938 | 0.64206 | 1.00000 | 0.98152 |
| NC_045512 | ORF10 | 0.66535 | 0.53126 | 0.62087 | 0.52204 | 0.88024 | 0.67895 | 0.94183 | 0.64000 | 0.65199 | 0.99573 | 0.97648 |
| MT012098 | ORF10 | 0.66535 | 0.53126 | 0.62087 | 0.52204 | 0.88024 | 0.67895 | 0.94183 | 0.64000 | 0.65199 | 0.99573 | 0.97903 |
| MT050493 | ORF10 | 0.66535 | 0.53126 | 0.62087 | 0.52204 | 0.88024 | 0.67895 | 0.94183 | 0.64000 | 0.65199 | 0.99573 | 0.97903 |
| MT358637 | ORF10 | 0.66535 | 0.53126 | 0.62087 | 0.52204 | 0.88024 | 0.67895 | 0.94183 | 0.64000 | 0.65199 | 0.99573 | 0.98152 |
| NC_045512 | ORF3a | 0.65733 | 0.60293 | 0.64026 | 0.54415 | 0.84395 | 0.74176 | 0.91830 | 0.69043 | 0.65731 | 0.99454 | 0.97677 |
| MT012098 | ORF3a | 0.65733 | 0.60293 | 0.64026 | 0.54415 | 0.84395 | 0.74176 | 0.91830 | 0.69043 | 0.65731 | 0.99454 | 0.97903 |
| MT050493 | ORF3a | 0.65733 | 0.60293 | 0.64026 | 0.54415 | 0.84395 | 0.74176 | 0.91830 | 0.69043 | 0.65731 | 0.99454 | 0.97903 |
| MT358637 | ORF3a | 0.65733 | 0.60293 | 0.64026 | 0.54415 | 0.84395 | 0.74176 | 0.91830 | 0.69043 | 0.65731 | 0.99454 | 0.98152 |
| NC_045512 | ORF6 | 0.64357 | 0.51174 | 0.61783 | 0.64681 | 0.94723 | 0.58368 | 0.93832 | 0.58368 | 0.60386 | 0.99992 | 0.97703 |
| MT012098 | ORF6 | 0.64357 | 0.51174 | 0.61783 | 0.64681 | 0.94723 | 0.58368 | 0.93832 | 0.58368 | 0.60386 | 0.99992 | 0.97903 |
| MT050493 | ORF6 | 0.64357 | 0.51174 | 0.61783 | 0.64681 | 0.94723 | 0.58368 | 0.93832 | 0.58368 | 0.60386 | 0.99992 | 0.97903 |
| MT358637 | ORF6 | 0.64357 | 0.51174 | 0.61783 | 0.64681 | 0.94723 | 0.58368 | 0.93832 | 0.58368 | 0.60386 | 0.99992 | 0.98152 |
| NC_045512 | ORF7a | 0.59207 | 0.58179 | 0.57340 | 0.52244 | 0.87520 | 0.75251 | 0.90698 | 0.65002 | 0.60385 | 0.99577 | 0.97710 |
| MT012098 | ORF7a | 0.59207 | 0.58179 | 0.57340 | 0.52244 | 0.87520 | 0.75251 | 0.90698 | 0.65002 | 0.60385 | 0.99577 | 0.97903 |
| MT050493 | ORF7a | 0.59207 | 0.58179 | 0.57340 | 0.52244 | 0.87520 | 0.75251 | 0.90698 | 0.65002 | 0.60385 | 0.99577 | 0.97903 |
| MT358637 | ORF7a | 0.59207 | 0.58179 | 0.57340 | 0.52244 | 0.87520 | 0.75251 | 0.90698 | 0.65002 | 0.60385 | 0.99577 | 0.98152 |
| NC_045512 | ORF8 | 0.56982 | 0.58887 | 0.63231 | 0.59225 | 0.84988 | 0.66871 | 0.94765 | 0.68673 | 0.59024 | 0.99515 | 0.97693 |
| MT012098 | ORF8 | 0.56982 | 0.58887 | 0.63231 | 0.59225 | 0.84988 | 0.66871 | 0.94765 | 0.68673 | 0.59024 | 0.99515 | 0.97903 |
| MT050493 | ORF8 | 0.56982 | 0.58962 | 0.62382 | 0.59225 | 0.84988 | 0.67479 | 0.94546 | 0.68673 | 0.59024 | 0.99515 | 0.97903 |
| MT358637 | ORF8 | 0.56982 | 0.58887 | 0.63231 | 0.59225 | 0.84988 | 0.66871 | 0.94765 | 0.68673 | 0.59024 | 0.99515 | 0.98152 |
| NC_045512 | ORF7b | 0.71222 | 0.54143 | 0.72032 | 0.52686 | 0.78637 | 0.68404 | 0.99403 | 0.55410 | 0.71541 | 0.94566 | 0.91796 |
| MT358637 | ORF7b | 0.71222 | 0.54143 | 0.72032 | 0.52686 | 0.78637 | 0.68404 | 0.99403 | 0.55410 | 0.71541 | 0.94566 | 0.98152 |

#### 2.1.1. Findings and Discussions on the gene E

Here in Table 4, we present codon usages of the gene E across the all four genomes.

**Table 4:** Codon usages and their corresponding frequencies

| Codons used in E | Freq. |
| --- | --- |
| CTT, GTT | 7 |
| TCT,TAC,AAT | 4 |
| TTC, GTA | 3 |
| TTT, TTG, CTA, CTG, GTG, CCT, ACA, GCG, AAA & TGC | 2 |
| ATG, TTA, ATT, ATC, ATA, GTC, TCA, TCG, ACT & ACG | 1 |
| GCT, GCC, TAA, AAC, GAT, GAA, GAG, TGT, CGT, CGA, AGT, AGC, AGA & GGT | 1 |
| CTC, TCC, CCC, CCA, CCG, ACC, GCA, TAT, TAG & CAT | 0 |
| CAC, CAA, CAG, AAG, GAC, TGA, TGG, CGC, CGG, AGG, GGC, GGA & GGG | 0 |

##### Codon usages

Table-4 shows that out of the six possibilities of the codons which code for L amino acid, only CTT has been chosen seven times in the protein sequence of E. The protein sequence of E contain the three amino acids say S, Y and N with frequency 4, which are coded by codons TCT, TAC and AAT respectively. This shows a simple codon biases over the gene E across all the four genomes. In contrast, it is found that the amino acid V is encoded in the primary protein sequence by three different codons such as GTA, GTC and GTT are used with different frequency viz. 3, 1, 7 respectively. Interestingly, amino acid Tryptophan (W) which is encoded by TGG only, does not appear in the protein sequence E over the four genomes. Similar to the codon TGG, there are several other codons which also do not appear in the gene E across the genomes.

##### Double nucleotide usages

The frequency of usage of the double nucleotides in the gene E over the four genomes are given in the Table 5.

**Table 5:** Frequency of double nucleotides over the gene E across the four genomes

| DN | Freq | DN | Freq |
| --- | --- | --- | --- |
| TT | 16 | TC | 7 |
| CT | 14 | AA | 6 |
| GT | 14 | CG | 5 |
| TA | 9 | GA | 4 |
| AT | 8 | GC | 4 |
| AC | 8 | CC | 3 |
| TG | 8 | CA | 1 |
| AG | 7 | GG | 0 |

The double nucleotide GG is not used at all in the gene E over the said genomes. It seems there is no bias of using double nucleotides unlike in the case of codon usages.

##### HEs and SEs of spatial representations

- From the Table 3, it follows that all the spatial representations of the nucleotides A, C, T and G of the gene E are positively trending in gene E over the four genomes. Based on the HEs obtained the positive trend of the bases A, C, T and G can be ordered as A, G, T and C. Among the four spatial representations, the most positively trending representation is of the nucleotide A which suggests that purine bases are positively trendier than pyrimidine bases.
- Further, SEs of spatial representation of the nucleotide T is highest compared to the other nucleotides.
- Table 3 follows that the nucleotide conservation entropy is turned out to be significant close to 1 for the gene E of the three genomes except the genome MT358637. The conservation entropy of the gene E in the genomes MT012098, MT050493 are extremely close whereas that of the gene E in the genomes NC_045512 is little different. This feature essentially discovers the conservation of A, T, C and G in the gene E of the genomes NC_045512, MT012098 are different though the sequential similarity of the genomes is 99.98%.

#### 2.1.2. Findings and Discussions on the gene M

##### Codon Usages

The frequency of codon usages in gene M over the four genomes has been elaborated in the Table 6.

**Table 6:**
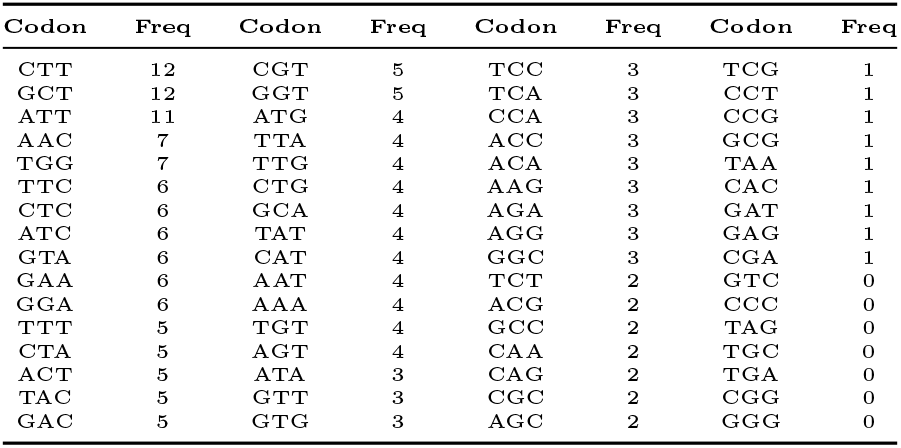
Frequency of usages over the gene M across the four genomes

It shows that the start codon ATG and stop codon TAA are with the frequencies four and once respectively in the gene M over the four genomes. Among the six codons which encode the amio acid L, the codon CTT is used in the gene M with highest frequency (12) suggesting the choice of codon bias. On the other side, all the three codons ATT, ATC and ATA which encode the amino acid (I) are present in the gene M with different frequency of usages. This shows there is no choice bias in selecting the codons in encoding (I) in the gene M. It is noted that the only codon TGG which encodes the amino acid Tryptophan (W) is used seven times in the gene M over the four genomes.

##### Double Nucleotide Usages

The frequencies of usage of double nucleotides over the gene M across the four genomes are illustrated in the Table 7. It follows that all possible double nucleotides are used in the gene M which indicates no choice bias.

**Table 7:** Frequency of double nucleotides over the gene M across the genomes

| DN | Freq | DN | Freq |
| --- | --- | --- | --- |
| TT | 32 | CA | 21 |
| TG | 30 | GT | 18 |
| TC | 29 | GC | 18 |
| TA | 27 | GG | 17 |
| AA | 25 | GA | 16 |
| CT | 23 | AG | 13 |
| AT | 22 | CC | 12 |
| AC | 21 | CG | 10 |

The double nucleotide TT is used maximum with frequency 32. It is worth noting that the unused double nucleotide GG in the gene E, is used seventeen times in the gene M of the four genomes.

##### HEs and SEs of spatial representations

- All the spatial representation of four nucleotides are positively trending over the gene Macross the four genomes. The order of nucleotides based on their respective trending behaviour over the gene is turned up as *T* > *A* > *C* > *G*. The HE of the purine-pyrimidine representation in the gene M over the genomes is 0.63095549 which signifies that the representations is positively trending.
- The SE of the binary representation of purine-pyrimidine bases over the gene E in the four genomes is 0.9961 which implies the amount of uncertainty is at almost maximum, i.e. the spatial arrangement of purine and pyrimidine bases are equally probable in the sequence of the gene M over all the four genomes. As in the case of the gene E, the amount of uncertainty of spatial arrangement of the nucleotide bases is highest for T.
- The nucleotide conservation entropy of the gene M over the genomes are getting varied from one to another and very close to 1 (maximum) as followed from Table 3. The conservation of nucleotide bases are different in respective spatial representations. The conservation entropies of nucleotides over the genomes NC_045512 and MT358637 are not as close as in MT012098 and MT050493 in the gene M. The minute difference among the M genes over all the four genomes is reflected through the conservation of the nucleotide bases, as observed in the case of gene E.

#### 2.1.3. Findings and Discussions on the gene N

##### Codon Usages

The frequency of codon usages in the gene N over the four genomes are presented in the Table 8. The frequency of usages are strictly same in three genomes NC_045512 (*G*_1_), MT012098 (*G*_2_) and MT050493 (*G*_3_) compared with MT358637 (G4) which is slightly varied. Note that codons TAG, TGT, TGC and TGA are absent in the gene N across the four genomes.

**Table 8:** Frequency of usages of codons over the gene M across the four genomes

| Codon | $G_1$ | $G_2$ | $G_3$ | $G_4$ | Codon | $G_1$ | $G_2$ | $G_3$ | $G_4$ |
| --- | --- | --- | --- | --- | --- | --- | --- | --- | --- |
| ATG | 7 | 7 | 7 | 7 | TAT | 2 | 2 | 2 | 2 |
| TTT | 3 | 3 | 3 | 3 | TAC | 9 | 9 | 9 | 9 |
| TTC | 10 | 10 | 10 | 10 | TAA | 1 | 1 | 1 | 1 |
| TTA | 2 | 2 | 2 | 2 | TAG | 0 | 0 | 0 | 0 |
| TTG | 9 | 9 | 9 | 9 | CAT | 3 | 3 | 3 | 3 |
| CTT | 8 | 8 | 8 | 8 | CAC | 1 | 1 | 1 | 1 |
| CTC | 2 | 2 | 2 | 2 | CAA | 27 | 27 | 27 | 27 |
| CTA | 3 | 3 | 3 | 3 | CAG | 8 | 8 | 8 | 8 |
| CTG | 3 | 3 | 3 | 3 | AAT | 16 | 16 | 16 | 16 |
| ATT | 9 | 9 | 9 | 10 | AAC | 6 | 6 | 6 | 6 |
| ATC | 4 | 4 | 4 | 4 | AAA | 21 | 21 | 21 | 21 |
| ATA | 1 | 1 | 1 | 1 | AAG | 10 | 10 | 10 | 10 |
| GTT | 2 | 2 | 2 | 2 | GAT | 14 | 14 | 14 | 14 |
| GTC | 3 | 3 | 3 | 3 | GAC | 10 | 10 | 10 | 10 |
| GTA | 1 | 1 | 1 | 1 | GAA | 8 | 8 | 8 | 8 |
| GTG | 2 | 2 | 2 | 2 | GAG | 4 | 4 | 4 | 4 |
| TCT | 8 | 8 | 8 | 8 | TGT | 0 | 0 | 0 | 0 |
| TCC | 3 | 3 | 3 | 3 | TGC | 0 | 0 | 0 | 0 |
| TCA | 9 | 9 | 9 | 9 | TGA | 0 | 0 | 0 | 0 |
| TCG | 2 | 2 | 2 | 2 | TGG | 5 | 5 | 5 | 5 |
| CCT | 8 | 8 | 8 | 8 | CGT | 6 | 6 | 6 | 6 |
| CCC | 7 | 7 | 7 | 7 | CGC | 5 | 5 | 5 | 5 |
| CCA | 11 | 11 | 11 | 11 | CGA | 5 | 5 | 5 | 6 |
| CCG | 2 | 2 | 2 | 2 | CGG | 2 | 2 | 2 | 1 |
| ACT | 16 | 16 | 16 | 15 | AGT | 9 | 9 | 9 | 9 |
| ACC | 6 | 6 | 6 | 6 | AGC | 6 | 6 | 6 | 6 |
| ACA | 8 | 8 | 8 | 8 | AGA | 10 | 10 | 10 | 10 |
| ACG | 2 | 2 | 2 | 2 | AGG | 1 | 1 | 1 | 1 |
| GCT | 19 | 19 | 19 | 19 | GGT | 10 | 10 | 10 | 10 |
| GCC | 7 | 7 | 7 | 7 | GGC | 16 | 16 | 16 | 16 |
| GCA | 8 | 8 | 8 | 8 | GGA | 13 | 13 | 13 | 13 |
| GCG | 3 | 3 | 3 | 3 | GGG | 4 | 4 | 4 | 4 |

The codon CAA and CAG, which encode the amino acid Glutamine (Q), are used respectively 27 and 8 times in the gene N. So there is no choice bias in selecting the codons for encoding the amino acid (Q). The codons CGT, CGC, CGA, CGG, AGA and AGG (encode the amino acid Arginine(R)) are all used in the gene N over the four genomes. No choice bias is seen in selecting the codons which encode Ariginine, Leuicine or Serine. Note that the codon TGG is used five times in the gene N over the four genomes. The only stop codon which is used in the gene N once is TAA.

##### Double Nucleotide Usages

The frequency usages of the double nucleotides is given in Table 9. Note that the frequency of the double nucleotides AT and GC in the gene N of the genome MT358637 is different from other three genomes as shown in Table 9.

**Table 9:**
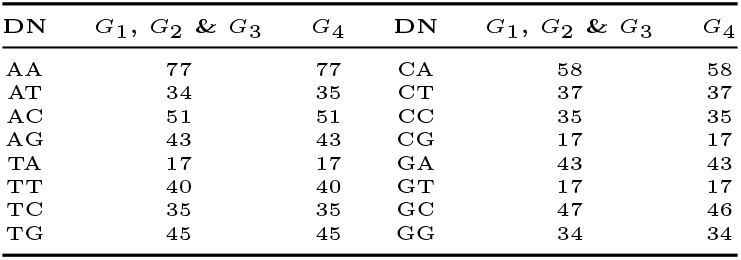
Frequency of double nucleotides over the gene N across the four genomes

| DN | $G_1, G_2 \& G_3$ | $G_4$ | DN | $G_1, G_2 \& G_3$ | $G_4$ |
| --- | --- | --- | --- | --- | --- |
| AA | 77 | 77 | CA | 58 | 58 |
| AT | 34 | 35 | CT | 37 | 37 |
| AC | 51 | 51 | CC | 35 | 35 |
| AG | 43 | 43 | CG | 17 | 17 |
| TA | 17 | 17 | GA | 43 | 43 |
| TT | 40 | 40 | GT | 17 | 17 |
| TC | 35 | 35 | GC | 47 | 46 |
| TG | 45 | 45 | GG | 34 | 34 |

It is noted that the highest frequency of usage is attained by the double nucleotide AA unlike in the previous cases. Clearly there is no bias of choices of double nucleotides in the gene N.

##### HEs and SEs of spatial representations

- Table 3 shows that the HE of the spatial representations of the nucleotides A, C, T and G in the gene N over the three genomes NC_045512, MT012098 and MT050493 are 0.61742, 0.50535, 0.53218 and 0.55515 respectively whereas that of the gene N over the genome MT358637 is turned out to be slightly different and they are 0.61484, 0.51273, 0.53644 and 0.55913 respectively for the A, C, T and G spatial representations. The most positively trending spatial representations is of the nucleotide A. The spatial representations of C is nearly random as the HE is turned out to be very close to 0.5. The binary spatial representation of the purine and pyrimidine bases over the gene N over the four genomes is surprisingly invariant and that is 0.57043. Clearly the spatial representation of purine and pyrimidine bases is positively trending.
- From the Table 3, it is observed that the SE of the spatial representations of the nucleotides A, C, T and G in the gene N over the three genomes NC_045512, MT012098 and MT050493 are 0.90160, 0.81128, 0.74209 and 0.90262 respectively. The SE of the binary representations of the nucleotides A, C, T and G in the gene N of the genome MT358637 are 0.90247, 0.81001, 0.74360 and 0.90262 respectively. The uncertainty of presence and absence of the purine and pyrimidine bases in the gene N over the four genomes is almost close to 1, implying that presence of purine and pyrimidine bases are equally likely over the gene N across the genomes.
- Note that the highest amount of uncertainty is present in the conservation of the nucleotide bases over the genome MT358637. Clearly, the uncertainty of nucleotides conservation in the gene N is getting varied from one to another in a very minute scale though the SEs of the gene N over the genomes MT012098, MT050493 based in Kerala are extremely nearer to each other.

#### 2.1.4. Findings and Discussions on the gene S

##### Codon usages

Here we list all the codons with their respective frequencies in the gene S over the four genomes NC_045512 (*G*_1_), MT012098(*G*_2_), MT050493(*G*_3_) and MT358637 (*G*_4_) in the Table 10.

**Table 10:** Frequency of codon usages in the gene S over the four genomes

| Codon | $G_1$ | $G_2$ | $G_3$ | $G_4$ | Codon | $G_1$ | $G_2$ | $G_3$ | $G_4$ |
| --- | --- | --- | --- | --- | --- | --- | --- | --- | --- |
| CCG | 0 | 0 | 0 | 0 | CAG | 16 | 16 | 16 | 16 |
| TAG | 0 | 0 | 0 | 0 | AGT | 17 | 17 | 17 | 17 |
| TGA | 0 | 0 | 0 | 0 | GGA | 17 | 17 | 17 | 17 |
| CGA | 0 | 0 | 0 | 1 | TTC | 18 | 18 | 18 | 18 |
| TAA | 1 | 1 | 1 | 1 | ATA | 18 | 19 | 18 | 18 |
| CGC | 1 | 1 | 1 | 1 | GAC | 19 | 19 | 19 | 19 |
| TCG | 2 | 2 | 2 | 2 | TTG | 20 | 20 | 20 | 20 |
| GCG | 2 | 2 | 2 | 2 | AGA | 20 | 19 | 20 | 20 |
| CGG | 2 | 2 | 2 | 2 | GTC | 21 | 21 | 21 | 21 |
| CTG | 3 | 3 | 3 | 3 | AAG | 23 | 23 | 23 | 23 |
| ACG | 3 | 3 | 3 | 3 | CCA | 25 | 25 | 25 | 25 |
| GGG | 3 | 3 | 3 | 3 | TCA | 26 | 26 | 26 | 26 |
| CCC | 4 | 4 | 4 | 4 | GCA | 27 | 27 | 27 | 27 |
| CAC | 4 | 4 | 4 | 4 | TTA | 28 | 28 | 28 | 28 |
| AGC | 5 | 5 | 5 | 5 | TGT | 28 | 28 | 28 | 28 |
| GCC | 8 | 8 | 8 | 8 | CCT | 29 | 29 | 29 | 29 |
| CTA | 9 | 9 | 9 | 9 | AAC | 34 | 34 | 34 | 34 |
| CGT | 9 | 9 | 9 | 9 | GAA | 34 | 34 | 34 | 34 |
| ACC | 10 | 10 | 10 | 10 | CTT | 36 | 36 | 36 | 36 |
| AGG | 10 | 10 | 10 | 10 | TCT | 37 | 37 | 37 | 37 |
| CTC | 12 | 12 | 12 | 12 | AAA | 38 | 38 | 38 | 38 |
| TCC | 12 | 12 | 12 | 12 | ACA | 40 | 40 | 40 | 40 |
| TGC | 12 | 12 | 12 | 12 | TAT | 40 | 39 | 40 | 40 |
| TGG | 12 | 12 | 12 | 12 | GCT | 42 | 42 | 41 | 42 |
| GTG | 13 | 13 | 13 | 13 | GAT | 43 | 43 | 43 | 42 |
| CAT | 13 | 13 | 13 | 13 | ATT | 44 | 44 | 44 | 44 |
| ATG | 14 | 14 | 14 | 14 | ACT | 44 | 44 | 44 | 44 |
| ATC | 14 | 14 | 14 | 14 | CAA | 46 | 46 | 46 | 45 |
| TAC | 14 | 14 | 14 | 14 | GGT | 47 | 47 | 47 | 48 |
| GAG | 14 | 14 | 14 | 14 | GTT | 48 | 48 | 49 | 48 |
| GTA | 15 | 15 | 15 | 15 | AAT | 54 | 54 | 54 | 54 |
| GGC | 15 | 15 | 15 | 15 | TTT | 59 | 59 | 59 | 59 |

The frequencies of codons such as ATA, AGA, TAT, GCT, GAT, CAA, GGT and GTT are perturbed over the four genomes. Note that the only codons TTT and TTC which encode the amino acid Phenylalanine (F) are present in the gene S with frequencies 59 and 18 respectively, suggesting no choice bias in selecting the codon for Phenylalanine. On contrary, the only stop codon TAA is present once in the gene S over the four genomes. The six codons TTA, TTG, CTT, CTC, CTA, CTG encoding Leucine(L) are present over the gene S in the four genomes with various frequencies. All the six possible codons, which encode the amino acid Arginine (R), are present in the gene M across all the four genomes. The codon TGG which encodes the amino acid (W), is present in the gene S over the four genomes with frequency 12. The codon CCG is never used in the gene S but the other codons which encode Proline(P) are all present in the S gene across the four genomes. Hence a slight choice bias exists in the gene S.

##### Double nucleotide usages

The double nucleotide frequency usages over the gene S of the four genomes are presented in the following Table 11.

**Table 11:** Double nucleotides frequencies in the gene S over the four genomes

| DN | $G_1$ | $G_2$ | $G_3$ | $G_4$ | DN | $G_1$ | $G_2$ | $G_3$ | $G_4$ |
| --- | --- | --- | --- | --- | --- | --- | --- | --- | --- |
| CG | 15 | 13 | 15 | 16 | CT | 116 | 144 | 115 | 116 |
| GC | 61 | 78 | 61 | 61 | AC | 118 | 131 | 118 | 118 |
| CC | 71 | 63 | 71 | 71 | AT | 138 | 164 | 138 | 137 |
| GG | 75 | 68 | 75 | 75 | TA | 139 | 124 | 139 | 139 |
| GA | 90 | 96 | 90 | 90 | CA | 155 | 141 | 155 | 154 |
| AG | 107 | 97 | 107 | 107 | TG | 166 | 166 | 166 | 166 |
| GT | 114 | 116 | 114 | 115 | AA | 189 | 185 | 189 | 189 |
| TC | 116 | 90 | 116 | 116 | TT | 241 | 233 | 242 | 241 |

It is found that all the sixteen double nucleotide are used in the S gene over the four genomes. The TT is used with highest frequency and CG is present with lowest frequency over the gene S of the four genomes. It is noticed that the only double nucleotide TG is present in the gene S over the four genomes, with equal frequency (166).

##### HEs and SEs of spatial representations

- From the Table 3, it is seen that all the spatial representations of the nucleotides as well as the purine-pyrimidine bases are positively trending as the HEs are coming out to be greater than 0.5. The HEs of each binary representations of A, T, C and G and in the purine-pyrimidine level of the gene S over the two genomes NC_045512 and MT050493 are almost same. The most positively trendiest spatial arrangement, of the nucleotide in the gene S over the four genomes, is base T as observed from the Table 3. It is noted that the HE of the spatial binary representation of the purine and pyrimidine bases over the gene S of the genomes are identical although the individual spatial representations of the nucleotides are not identical.
- The amount of uncertainty of occurrence of purine and pyrimidine bases over the gene S of the four genomes is at maximum and that implies the equal probability of occurrence of the purine and pyrimidine bases in the gene S. Also the SEs are turned out to be same for the spatial binary representations of the purine-pyrimidine bases.
- The SE of the nucleotide conservation entropy of the gene S over the four genomes NC_045512, MT012098, MT050493 and MT358637 are 0.9764, 0.9790, 0.9790 and 0.9815 respectively. The amount of uncertainty of conservation of nucleotides is high in the gene S over the four genomes. It is observed that the conservation of nucleotides are very much close for the gene S in the genomes MT012098 and MT050493.

#### 2.1.5. Findings and Discussions on the gene ORF1

The longest gene ORF1 which encodes 16 non-structural proteins of length 21290 across the four genomes. Although the length is fixed among all the genomes, there is a slight frequency variance of the nucleotide bases as presented in the Table 2. Here in addition, the nucleotide density and AT-GC density of the gene ORF1 across the four genomes are figured in the Fig.3.

**Figure 3:**
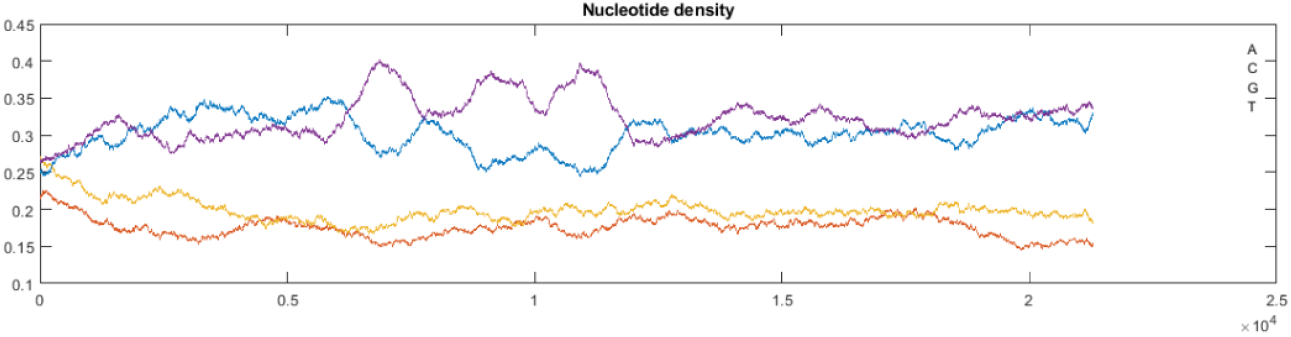
Nucleotide density and AT-GC density over the gene ORF1 across the four genomes

Fig.3 follows that the density of the bases A (30.18%) and T (32.18%) in the gene ORF1 across the four genomes dominate the density of the other two bases C (17.59%) and G (19.87%) as depicted in the Fig.3.

##### Codon usages

The frequency of codon usages with its graphical representations (Fig.4) in the gene ORF1 over the four genomes are given in the Table 12. It is found that all the sixty four codons are used over the gene ORF1 across the four genomes. The frequency of usages of codons in the gene ORF1 across the four genomes is very much robust, that is, the amount of variation of frequency from one genome to another is very small as observed. The start codon ATG attains the highest frequency which is 266 in the gene ORF1 across four genomes. It is noticed that all the three stop codons TAG, TGA and TAA are used with varied frequencies 72, 42 and 58 respectively.

**Table 12:**
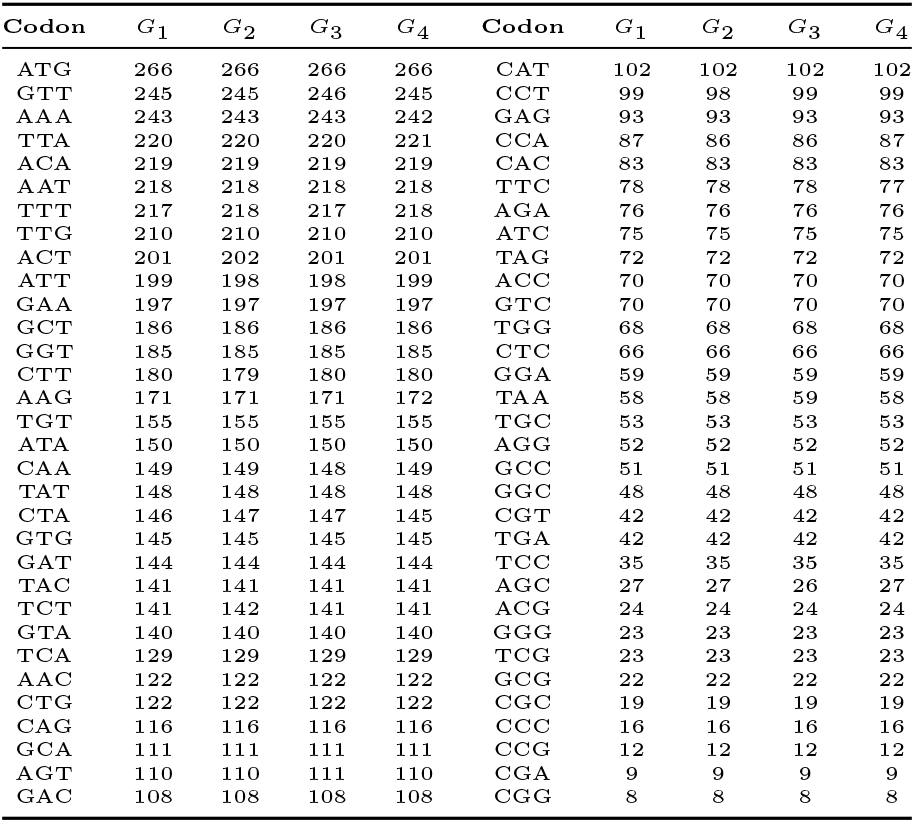
frequency of codon usages in the gene ORF1 over the four genomes

**Figure 4:**
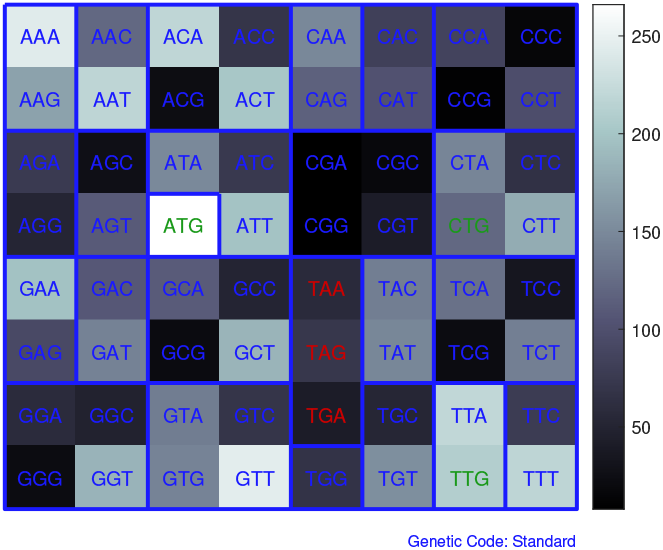
Graphical representation of usages of codons over the gene ORF1 across the four genomes

Clearly, there is no codon bias of choices in the gene ORF1 over the four genomes as shown in the Fig.5.

**Figure 5:**
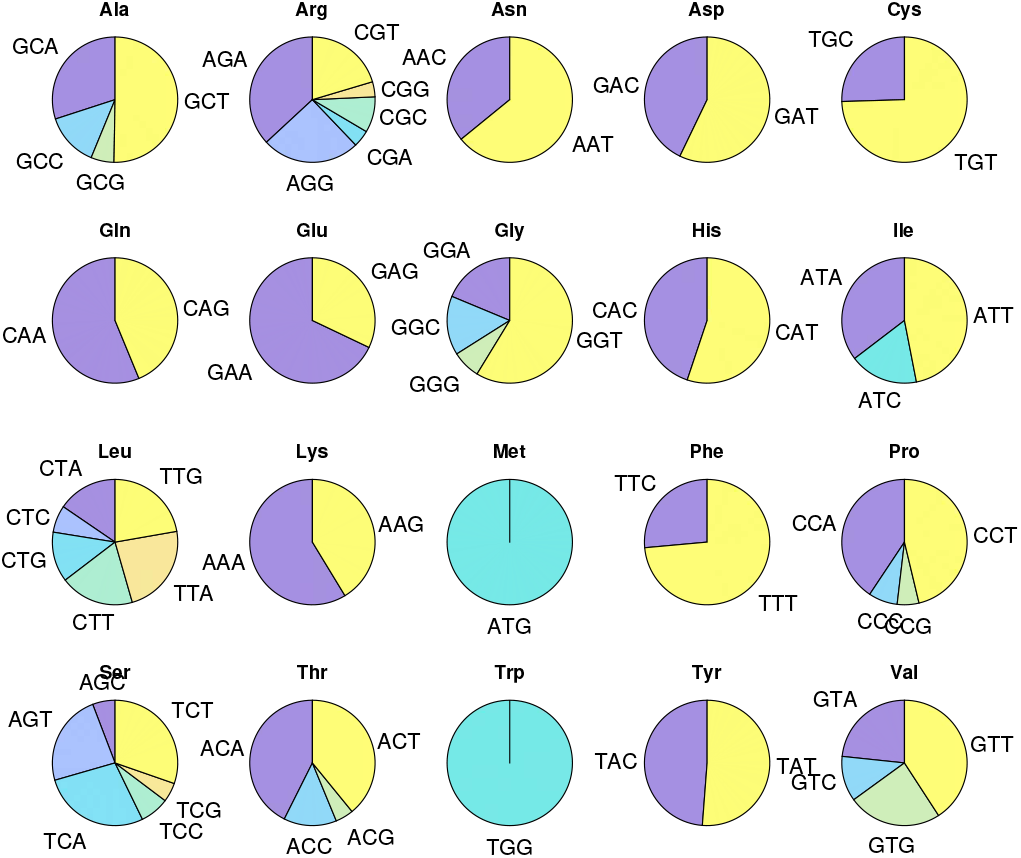
Codon usages based on amino acids over the gene ORF1 across the four genomes

Following in the Table 13, the frequency of amino acids in the primary protein sequence of ORF1 with graphical frequency distribution as shown in Fig.6.

**Table 13:** Frequencies of amino acids over the protein sequence of ORF1

| AA | Freq | AA | Freq |
| --- | --- | --- | --- |
| L | 944 | E | 290 |
| V | 600 | Y | 289 |
| T | 514 | M | 266 |
| I | 424 | Q | 265 |
| K | 414 | D | 252 |
| A | 370 | P | 214 |
| N | 340 | C | 208 |
| S | 336 | R | 206 |
| G | 315 | H | 185 |
| F | 295 | STOP | 172 |
|  |  | W | 68 |

**Figure 6:**
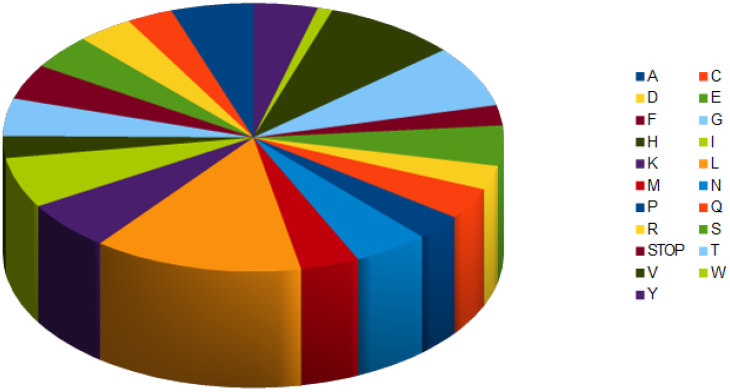
Pie chart of frequency distribution of amino acids over the gene ORF1

The Table 13 follows that the amino acid Leucine is maximally used with frequency 944 over the protein ORF1 whereas Tryptophan is least used in ORF1.

##### Double nucleotide usages

The frequencies of double nucleotides over the gene ORF1 of the four genomes are presented in the Table 14 with bar plot in the Fig.7. It is seen that all the sixteen double nucleotides are present over the gene ORF1 across the four genomes.

**Table 14:** frequencies of double nucleotides over the gene ORF1 of the four genomes

| DN | $G_1$ | $G_2$ | $G_3$ | $G_4$ | DN | $G_1$ | $G_2$ | $G_3$ | $G_4$ |
| --- | --- | --- | --- | --- | --- | --- | --- | --- | --- |
| AA | 1004 | 1004 | 1004 | 1003 | CA | 722 | 722 | 722 | 722 |
| AT | 869 | 868 | 870 | 869 | CT | 734 | 735 | 735 | 733 |
| AC | 717 | 718 | 716 | 717 | CC | 302 | 301 | 300 | 302 |
| AG | 609 | 609 | 609 | 609 | CG | 149 | 149 | 149 | 149 |
| TA | 914 | 914 | 913 | 914 | GA | 586 | 586 | 586 | 587 |
| TT | 1133 | 1133 | 1133 | 1135 | GT | 756 | 758 | 756 | 756 |
| TC | 415 | 415 | 416 | 414 | GC | 403 | 401 | 403 | 403 |
| TG | 937 | 937 | 938 | 937 | GG | 395 | 395 | 395 | 395 |

**Figure 7:**
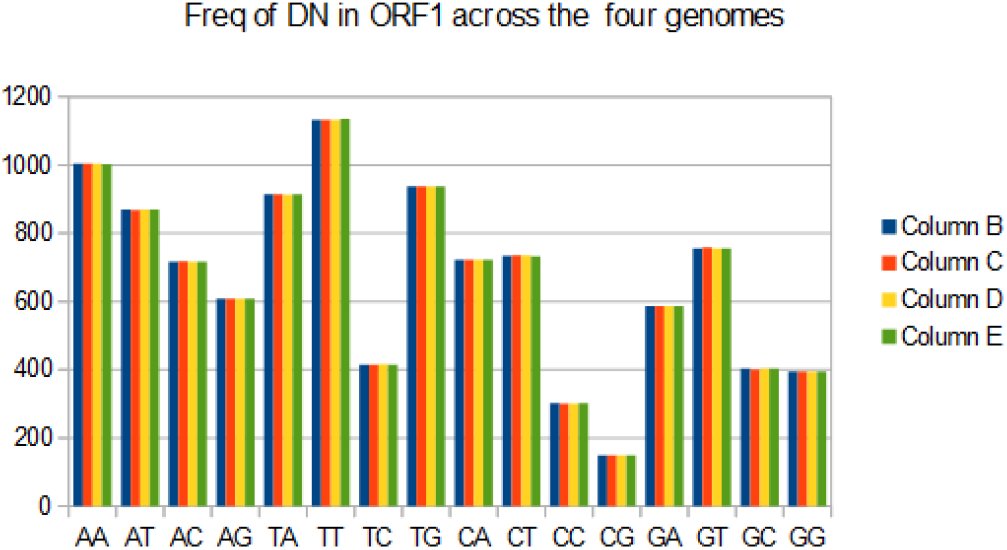
Bar plot of the double nucleotide usages over the gene ORF1 across the four genomes. Here the columns B, C, D, E denote in the figure the genomes *G*_1_, *G*_2_, *G*_3_ and *G*_4_ respectively.

It is observed that the most frequently used double nucleotides is TT with frequency 1133 in ORF1 across all the four genomes. It is noteworthy that the frequency of TT in ORF1 over the Indian genome MT358637 is 1135. There is not much variance of frequencies of double nucleotides in ORF1 from one genome to another, is observed.

##### HEs and SEs of spatial representations

- From the Table 3, it is quite clear that the spatial representations are positively trending. Each of the four nucleotide spatial representation has its own positive autocorrelation (trending behaviour) as the HEs are different significantly from one to another. The decreasing order of positive trendiness of each nucleotide is *T > A > G > C*. The spatial organization of the purine-pyrimidine bases of the gene ORF1 is positively trending identically across the four genomes.
- The SE of each of the spatial representations of nucleotides are almost identical of the gene ORF1 across the genomes. The uncertainty of the presence of purine bases over the purine-pyrimidine representation of the gene ORF1 over the four genomes is at maximum.
- From the SE of the nucleotide conservations as presented in the Table 3, it is observed that the conservation of nucleotides over the gene ORF1 of the genomes MT012098 and MT050493 are close enough. The highest amount of uncertainty of information of conservations of nucleotides appears in the gene ORF1 over the genome NC_045512.

#### 2.1.6. Findings and Discussions on the gene ORF10

##### Codon usages

The frequency of usages of codons in the gene ORF10 are given in the Table 15. Due to the absence of the necessary codons, the amino acids Tryptophan and Glutamine are absent in the protein sequence of ORF10. It emerged that among three possible stop codons, the only stop codon TAG is present in ORF10.

**Table 15:** Codon usages and its frequencies over the gene ORF10 across four genomes.

| Codon | Freq | Codon | Freq | Codon | Freq | Codon | Freq |
| --- | --- | --- | --- | --- | --- | --- | --- |
| ATG | 2 | TCT | 1 | TAT | 2 | TGT | 0 |
| TTT | 3 | TCC | 0 | TAC | 1 | TGC | 1 |
| TTC | 1 | TCA | 0 | TAA | 0 | TGA | 0 |
| TTA | 0 | TCG | 0 | TAG | 1 | TGG | 0 |
| TTG | 1 | CCT | 0 | CAT | 0 | CGT | 1 |
| CTT | 0 | CCC | 0 | CAC | 0 | CGC | 0 |
| CTC | 2 | CCA | 0 | CAA | 1 | CGA | 0 |
| CTA | 1 | CCG | 1 | CAG | 0 | CGG | 0 |
| CTG | 0 | ACT | 0 | AAT | 2 | AGT | 1 |
| ATT | 0 | ACC | 0 | AAC | 3 | AGC | 0 |
| ATC | 0 | ACA | 1 | AAA | 0 | AGA | 1 |
| ATA | 3 | ACG | 1 | AAG | 0 | AGG | 0 |
| GTT | 2 | GCT | 1 | GAT | 1 | GGT | 0 |
| GTC | 0 | GCC | 0 | GAC | 0 | GGC | 1 |
| GTA | 2 | GCA | 1 | GAA | 0 | GGA | 0 |
| GTG | 0 | GCG | 0 | GAG | 0 | GGG | 0 |

Among the four possible codons, only the CCG has been chosen to encode the amino acid Proline(P) in the gene ORF10. Thus a choice bias codon is observed in the gene ORF10 across the four genomes.

##### Double nucleotide usages

The usages of double nucleotides over the gene ORF10 is tabulated in Table 16.

**Table 16:** Frequency of double nucleotides over the gene ORF10 across the four genomes

| DN | Freq | DN | Freq |
| --- | --- | --- | --- |
| TA | 10 | TG | 3 |
| AT | 8 | GA | 3 |
| TT | 5 | AA | 2 |
| CA | 5 | AC | 2 |
| CT | 5 | GT | 2 |
| TC | 4 | GC | 1 |
| CG | 4 | GG | 1 |
| AG | 3 | CC | 0 |

The double nucleotide CC is thoroughly absent from the gene ORF10 across the four genomes. It is noted that the TA is having highest frequency in the gene ORF10 unlike in the previous cases. It is worth noting that in the aforementioned genes double nucleotides AA and TT were having highest frequencies in the corresponding gene.

##### HEs and SEs of spatial representations

- From the HEs of genes ORF10 across four genomes, it is observed that the spatial representations are turned out to be positively auto-correlated. The spatial binary representation of the nucleotide A is the most auto-correlated representation whereas the least auto-correlated spatial representation is of the nucleotide G.
- As in the case of others genes, the uncertainty is at maximum of the presence and absence of pyrimidine bases in ORF10 across the four genomes.
- The SE of conservation of nucleotides over the gene ORF10 across the four genomes NC_045512, MT012098, MT050493 and MT358637 are turned out to be 0.97648, 0.97903, 0.97903 and 0.98152. This illustrates that the nucleotide conservations in the gene ORF10 over the genomes MT012098, MT050493 are identical. The uncertainty of conservation of nucleotides over the gene ORF10 is high as the SEs are closed to 1.

#### 2.1.7. Findings and Discussions on the gene ORF3a

##### Codon usages

The frequency of codon usages over the the gene ORF3a across the four genomes is presented in the Table 17. Note that that the codons TCG, CCC, GCG, TAG, TGA, CGA, CGG and GGG do not appear in the gene ORF3a.

**Table 17:** Frequency of codons in ORF3 across the four genomes

| Codon | Freq | Codon | Freq | Codon | Freq | Codon | Freq |
| --- | --- | --- | --- | --- | --- | --- | --- |
| ATG | 4 | TCT | 3 | TAT | 8 | TGT | 3 |
| TTT | 8 | TCC | 4 | TAC | 9 | TGC | 4 |
| TTC | 6 | TCA | 8 | TAA | 1 | TGA | 0 |
| TTA | 3 | TCG | 0 | TAG | 0 | TGG | 6 |
| TTG | 9 | CCT | 7 | CAT | 4 | CGT | 1 |
| CTT | 10 | CCC | 0 | CAC | 4 | CGC | 1 |
| CTC | 5 | CCA | 3 | CAA | 5 | CGA | 0 |
| CTA | 1 | CCG | 2 | CAG | 4 | CGG | 0 |
| CTG | 2 | ACT | 13 | AAT | 4 | AGT | 5 |
| ATT | 9 | ACC | 2 | AAC | 4 | AGC | 2 |
| ATC | 5 | ACA | 6 | AAA | 7 | AGA | 3 |
| ATA | 7 | ACG | 3 | AAG | 4 | AGG | 1 |
| GTT | 14 | GCT | 7 | GAT | 7 | GGT | 7 |
| GTC | 3 | GCC | 3 | GAC | 6 | GGC | 3 |
| GTA | 7 | GCA | 3 | GAA | 10 | GGA | 4 |
| GTG | 1 | GCG | 0 | GAG | 1 | GGG | 0 |

The stop codons TAG and TGA are not at all used in the gene ORF3a over the four genomes. All the necessary codons for encoding twenty amino acids are present in the gene ORF3a.

##### Double nucleotide usages

All the sixteen double nucleotides are present in the gene ORF3a across the four genomes as shown in Table 18.

**Table 18:** Frequency of usages of double nucleotides over the gene ORF3a across the genomes

| DN | Freq | DN | Freq |
| --- | --- | --- | --- |
| AA | 39 | CA | 29 |
| AT | 24 | CT | 38 |
| AC | 29 | CC | 11 |
| AG | 13 | CG | 7 |
| TA | 24 | GA | 28 |
| TT | 53 | GT | 26 |
| TC | 24 | GC | 25 |
| TG | 34 | GG | 10 |

The highest frequency is attained by the double nucleotide TT in ORF3a over the four genomes. So there is no choice bias of double nucleotides in ORF3a.

##### HEs and SEs of spatial representations

- The spatial representations of A, T, C and G as well as of the purine-pyrimidine bases are found to be positively auto-correlated.
- SEs of the binary representations of the nucleotides A, C, T and G over the gene ORF3a across the genomes are invariant as found in the Table 3. The SE of the binary spatial representation of the purine and pyrimidine bases of the gene ORF3a across the four genomes is 0.99454 which is very closed to 1 and that represents maximum uncertainty.
- From the nucleotide conservation SEs of ORF3a as found in the Table 3, it is found that the nucleotide conservations over ORF3a across the two genomes MT012098 and MT050493 are similar.

#### 2.1.8. Findings and Discussions on the gene ORF6

##### Codon usages

The frequency of each codon in ORF6 is remain invariant across the genomes as observed from the Table 19.

**Table 19:** Frequency of codon usages over the gene ORF6 across the four genomes

| Codon | Freq | Codon | Freq | Codon | Freq | Codon | Freq |
| --- | --- | --- | --- | --- | --- | --- | --- |
| ATG | 3 | TCT | 2 | TAT | 1 | TGT | 0 |
| TTT | 3 | TCC | 1 | TAC | 1 | TGC | 0 |
| TTC | 0 | TCA | 1 | TAA | 1 | TGA | 0 |
| TTA | 3 | TCG | 0 | TAG | 0 | TGG | 1 |
| TTG | 0 | CCT | 0 | CAT | 1 | CGT | 0 |
| CTT | 1 | CCC | 0 | CAC | 0 | CGC | 0 |
| CTC | 2 | CCA | 1 | CAA | 2 | CGA | 0 |
| CTA | 2 | CCG | 0 | CAG | 1 | CGG | 0 |
| CTG | 0 | ACT | 3 | AAT | 3 | AGT | 0 |
| ATT | 5 | ACC | 0 | AAC | 1 | AGC | 0 |
| ATC | 1 | ACA | 0 | AAA | 3 | AGA | 0 |
| ATA | 4 | ACG | 0 | AAG | 1 | AGG | 1 |
| GTT | 3 | GCT | 0 | GAT | 3 | GGT | 0 |
| GTC | 0 | GCC | 0 | GAC | 1 | GGC | 0 |
| GTA | 0 | GCA | 1 | GAA | 1 | GGA | 0 |
| GTG | 0 | GCG | 0 | GAG | 4 | GGG | 0 |

The interesting findings here: The only codon GCA out four codons, AGG out of six codons and CCA out of four codons which encode the amino acids Alanine (A), Arginine (R) and Prolin (P) respectively, are present in the gene ORF6 across the four genomes. Therefore there is a strong codon bias exists in ORF6 across the four genomes. The amino acids Cysteine and Glycine are absent from the protein sequence ORF6 as the necessary codons are absent in the gene sequence of ORF6.

##### Double nucleotide usages

The only double nucleotide CG is absent in the gene ORF6 thoroughly over the four genomes as presented in Table 20.

**Table 20:** Frequency of usages of double nucleotides over the gene ORF6 across the four genomes

| DN | Freq | DN | Freq |
| --- | --- | --- | --- |
| TA | 13 | CA | 6 |
| TT | 12 | AG | 5 |
| AA | 10 | CT | 4 |
| AT | 9 | GT | 3 |
| GA | 9 | CC | 1 |
| TC | 7 | GC | 1 |
| AC | 6 | GG | 1 |
| TG | 6 | CG | 0 |

The double nucleotide TA GG attain the highest and lowest frequency respectively in the gene ORF6 across the four genomes.

##### HEs and SEs of spatial representations

- Clearly from the HEs as observed in Table 3, all five different spatial representations (four nucleotides and purine-pyrimidine) are positively auto-correlated/trending in the gene ORF6 across the four genomes.
- From the Table 3, it is seen that the SEs of the representation of A and T are almost same and hence it is concluded that the spatial representations of A (purine-base) and T (pyrimidine-base) are similar. The SE of the spatial representation of purine and pyrimidine bases in ORF6 over the four genomes are again found same and that is 0.99992 which is very nearer to one.
- The SEs of the conservation of nucleotides over ORF6 across the three genomes NC_045512, MT012098, MT050493 is identical with the value 0.97703 whereas that in the genome MT358637 is 0.98152.

#### 2.1.9. Findings and Discussions on the gene ORF7a

##### Codon usages

The codons usages over the gene ORF7a across the four genomes is tabulated in the Table 21. It is observed that the frequencies of codon usages over the four genomes are turned out to be invariant.

**Table 21:** Frequency of codons usage in ORF7a across the four genomes

| Codon | Freq | Codon | Freq | Codon | Freq | Codon | Freq |
| --- | --- | --- | --- | --- | --- | --- | --- |
| TTT | 7 | TCA | 3 | ATC | 1 | GGA | 1 |
| ACA | 7 | ACT | 3 | GTC | 1 | TCC | 0 |
| CTT | 6 | GCA | 3 | GTG | 1 | TCG | 0 |
| AAA | 6 | TAC | 3 | GCC | 1 | CCC | 0 |
| ATT | 4 | TGT | 3 | GCG | 1 | CCG | 0 |
| GTT | 4 | TGC | 3 | CAT | 1 | ACC | 0 |
| CCT | 4 | TTA | 2 | CAG | 1 | ACG | 0 |
| GCT | 4 | CTC | 2 | AAT | 1 | TAA | 0 |
| CAA | 4 | GTA | 2 | AAC | 1 | TAG | 0 |
| GAA | 4 | CCA | 2 | AAG | 1 | TGG | 0 |
| GAG | 4 | TAT | 2 | GAT | 1 | CGC | 0 |
| AGA | 4 | CAC | 2 | GAC | 1 | CGA | 0 |
| TTC | 3 | GGC | 2 | TGA | 1 | CGG | 0 |
| CTG | 3 | ATG | 1 | CGT | 1 | AGT | 0 |
| ATA | 3 | TTG | 1 | AGC | 1 | AGG | 0 |
| TCT | 3 | CTA | 1 | GGT | 1 | GGG | 0 |

The amino acid Leusine(L) is encoded by six different amino acids which all are present with different frequencies over the gene ORF7a across the four genomes. Out of the six codons only two codon AGG and CGT have been chosen to encode Arginine (R) by the gene ORF7a. Hence a partial choice bias is observed. The amino acid (W) would not be present in ORF7a protein since the necessary codon TGG is absent in the sequence of ORF7a. It is noted that the one stop codon TGA is used once over the gene ORF7a across the four genomes.

##### Double nucleotide usages

The Table 22 follows that all the double nucleotides are present with atleast frequency greater than or equals to 2.

**Table 22:** Frequency of double nucleotides over the gene ORF7a across the genomes

| DN | Freq | DN | Freq |
| --- | --- | --- | --- |
| AA | 17 | CA | 11 |
| AT | 9 | CT | 18 |
| AC | 21 | CC | 2 |
| AG | 10 | CG | 2 |
| TA | 9 | GA | 14 |
| TT | 24 | GT | 5 |
| TC | 15 | GC | 8 |
| TG | 14 | GG | 4 |

The highest frequency 24 is attained by the double nucleotide TT as found in ORF7a. The double nucleotides CC and CG both are having least frequency which is 2.

##### HEs and SEs of spatial representations

- The spatial representation of A is found to be most positively trending as compared to others since the HE is maximum among all. The HE of the purine-pyrimidine arrangements is 0.60385 which depicts the positive auto-correlation over the representation.
- From the SEs of nucleotide bases in ORF7a as mentioned in Table 3, it is observed that the amount of uncertainty of presence of the nucleotide G over its binary representations is lowest as compared to others. The SE of binary representation of the purine-pyrimidine bases over the gene ORF7a across the genomes are also same which is 0.99577. As usual the highest amount of uncertainty is observed in the presence of purine bases over the gene ORF7a across the four genomes.
- The SE of conservation of nucleotides in the gene ORF7a over the four genomes NC_045512, MT012098, MT050493 and MT358637 are 0.97710, 0.97903, 0.97903 and 0.98152. The conservation of nucleotide bases are found to be similar in ORF7a over the two genomes MT012098, MT050493 as the SEs in these two cases are found to be identical. As previously seen, in the genome MT358637, the SE of nucleotide conservation in ORF7a is found to be more uncertain as the SE is seen to be close enough to 1.

#### 2.1.10. Findings and Discussions on the gene ORF8

##### Codon usages

The codon frequencies in the gene ORF8 across the four genomes are given in the following Table 23.

**Table 23:** Frequencies of codon usages in ORF8 across the four genomes.

| Codon | $G_1$ | $G_2$ | $G_3$ | $G_4$ | Codon | $G_1$ | $G_2$ | $G_3$ | $G_4$ |
| --- | --- | --- | --- | --- | --- | --- | --- | --- | --- |
| ATG | 1 | 1 | 1 | 1 | TAT | 6 | 6 | 6 | 6 |
| TTT | 4 | 4 | 4 | 4 | TAC | 1 | 1 | 1 | 1 |
| TTC | 4 | 4 | 4 | 4 | TAA | 1 | 1 | 1 | 1 |
| TTA | 6 | 6 | 5 | 6 | TAG | 0 | 0 | 0 | 0 |
| TTG | 2 | 2 | 2 | 2 | CAT | 2 | 2 | 2 | 2 |
| CTT | 2 | 2 | 2 | 2 | CAC | 2 | 2 | 2 | 2 |
| CTC | 0 | 0 | 0 | 0 | CAA | 3 | 3 | 3 | 3 |
| CTA | 0 | 0 | 0 | 0 | CAG | 3 | 3 | 3 | 3 |
| CTG | 0 | 0 | 0 | 0 | AAT | 2 | 2 | 2 | 2 |
| ATT | 5 | 5 | 5 | 5 | AAC | 0 | 0 | 0 | 0 |
| ATC | 5 | 5 | 5 | 5 | AAA | 5 | 5 | 5 | 5 |
| ATA | 0 | 0 | 0 | 0 | AAG | 0 | 0 | 0 | 0 |
| GTT | 6 | 6 | 6 | 6 | GAT | 4 | 4 | 4 | 4 |
| GTC | 0 | 0 | 0 | 0 | GAC | 3 | 3 | 3 | 3 |
| GTA | 4 | 4 | 4 | 4 | GAA | 4 | 4 | 4 | 4 |
| GTG | 2 | 2 | 2 | 2 | GAG | 2 | 2 | 2 | 2 |
| TCT | 2 | 2 | 2 | 2 | TGT | 5 | 5 | 5 | 5 |
| TCC | 1 | 1 | 1 | 1 | TGC | 2 | 2 | 2 | 2 |
| TCA | 3 | 3 | 4 | 3 | TGA | 0 | 0 | 0 | 0 |
| TCG | 1 | 1 | 1 | 1 | TGG | 1 | 1 | 1 | 1 |
| CCT | 4 | 4 | 4 | 4 | CGT | 2 | 2 | 2 | 2 |
| CCC | 1 | 1 | 1 | 1 | CGC | 0 | 0 | 0 | 0 |
| CCA | 1 | 1 | 1 | 1 | CGA | 0 | 0 | 0 | 0 |
| CCG | 1 | 1 | 1 | 1 | CGG | 0 | 0 | 0 | 0 |
| ACT | 2 | 2 | 2 | 2 | AGT | 2 | 2 | 2 | 2 |
| ACC | 0 | 0 | 0 | 0 | AGC | 0 | 0 | 0 | 0 |
| ACA | 3 | 3 | 3 | 3 | AGA | 2 | 2 | 2 | 2 |
| ACG | 0 | 0 | 0 | 0 | AGG | 0 | 0 | 0 | 0 |
| GCT | 3 | 3 | 3 | 3 | GGT | 3 | 3 | 3 | 3 |
| GCC | 0 | 0 | 0 | 0 | GGC | 0 | 0 | 0 | 0 |
| GCA | 2 | 2 | 2 | 2 | GGA | 2 | 2 | 2 | 2 |
| GCG | 0 | 0 | 0 | 0 | GGG | 0 | 0 | 0 | 0 |

The preferred stop codon in ORF8 is TAA across the four genomes and it is used once only. It is worth mentioning that the same length (366 bases) gene ORF7a uses only the stop codon TGA. Interestingly, all the twenty amino acid are present ORF8 protein.

##### Double nucleotide usages

A list of frequency of double nucleotides over the gene ORF8 across the four genomes is given in the Table 24. It is found that all the sixteen double nucleotides are used in ORF8 across the four genomes. The highest frequency of TT in the gene ORF8 is turned out to be 24.

**Table 24:** Frequency of double nucleotides over the gene ORF8 across the four genomes

| DN | $G_1$ | $G_2$ | $G_3$ | $G_4$ | DN | $G_1$ | $G_2$ | $G_3$ | $G_4$ |
| --- | --- | --- | --- | --- | --- | --- | --- | --- | --- |
| **GG** | 3 | 3 | 3 | 3 | **AA** | 13 | 13 | 13 | 13 |
| **CG** | 5 | 5 | 5 | 5 | **GA** | 13 | 13 | 13 | 13 |
| **GC** | 5 | 5 | 5 | 5 | **TG** | 14 | 14 | 14 | 14 |
| **CC** | 6 | 6 | 6 | 6 | **GT** | 14 | 14 | 14 | 14 |
| **AC** | 7 | 7 | 7 | 7 | **AT** | 16 | 16 | 16 | 16 |
| **CT** | 8 | 8 | 8 | 8 | **TC** | 16 | 16 | 16 | 16 |
| **AG** | 10 | 10 | 10 | 10 | **TA** | 18 | 18 | 17 | 18 |
| **CA** | 11 | 11 | 12 | 11 | **TT** | 24 | 24 | 24 | 24 |

It is found that the double nucleotide CA is present twelve times in the gene ORF8 over the genome MT050493 whereas in the other three genomes the gene ORF8 contains CA only eleven times uniformly. Also it is observed that TA is present in ORF8 over the genome MT050493 seventeen times whereas in the rests genomes it is present eighteen times in the gene ORF8.

##### HEs and SEs of spatial representations

- The highest amount of autocorrelation is observed in the spatial representation of T as found from the Table 3. The HE of the spatial representation of the purine and pyrimidine bases in ORF8 across the four genomes is 0.59024.
- The SEs of the binary representations of the nucleotides A, C, T and G in ORF8 over the three genomes except MT050493 are found to be same and they are 0.84988, 0.66871, 0.94765 and 0.68673 respectively whereas that in the gene ORF8 over the genome MT050493 are respectively 0.84988, 0.67479, 0.94546 and 0.68673. That is the the spatial template of the nucleotides C and T are different from that of the gene ORF8 over the other three genomes NC_045512, MT012098 and MT358637. As usual the binary SE of the spatial representation of the purine and pyrimidine bases over the gene ORF8 across the four genomes are found to be same and that is 0.99515.
- From the Table 3, it is found that the SEs of nucleotide conservations over the gene ORF8 across the three genomes except MT358637 is same as 0.97903 whereas that of the gene ORF8 in the genome MT358637 is 0.98152. As seen before the highest amount uncertainty is observed in conservation of nucleotides over the gene ORF8 of the genome MT358637. It is noted that the conservation of nucleotides of the gene ORF8 over the other three genomes are observed to be similar as the SEs are identical.

#### 2.1.11. Findings and Discussions on the gene ORF7b

From the Table 1, it is quite evident that the presence of ORF7b gene makes the two genomes NC_045512 and MT358637 separate from the rest two. It is worth noting that the absence and presence of ORF7b can be associated to the S and L type respectively. The features encountered for ORF7b over the two genomes NC_045512 and MT358637 are presented below:

##### Codon usages

The codon ATG (start codon), CTG, TCA and GCC are present twice over the gene ORF7b as observed in the Table 25. It is noted that the codon TAA is the most preferred stop codon among the three. There are 38 codons which are not present in the gene ORF7b in the two genomes.

**Table 25:** Frequency of codon usages over ORF7b in the two genomes

| Codon | Freq | Codon | Freq | Codon | Freq | Codon | Freq |
| --- | --- | --- | --- | --- | --- | --- | --- |
| ATG | 2 | TCT | 0 | TAT | 1 | TGT | 1 |
| TTT | 3 | TCC | 0 | TAC | 0 | TGC | 1 |
| TTC | 3 | TCA | 2 | TAA | 1 | TGA | 0 |
| TTA | 3 | TCG | 0 | TAG | 0 | TGG | 1 |
| TTG | 1 | CCT | 0 | CAT | 1 | CGT | 0 |
| CTT | 4 | CCC | 0 | CAC | 1 | CGC | 0 |
| CTC | 0 | CCA | 0 | CAA | 1 | CGA | 0 |
| CTA | 1 | CCG | 0 | CAG | 0 | CGG | 0 |
| CTG | 2 | ACT | 1 | AAT | 1 | AGT | 0 |
| ATT | 4 | ACC | 0 | AAC | 0 | AGC | 0 |
| ATC | 1 | ACA | 0 | AAA | 0 | AGA | 0 |
| ATA | 0 | ACG | 0 | AAG | 0 | AGG | 0 |
| GTT | 1 | GCT | 0 | GAT | 1 | GGT | 0 |
| GTC | 0 | GCC | 2 | GAC | 1 | GGC | 0 |
| GTA | 0 | GCA | 0 | GAA | 3 | GGA | 0 |
| GTG | 0 | GCG | 0 | GAG | 0 | GGG | 0 |

The amino acids Lysine, Arginine, Glycine and Proline are absent from the protein sequence of ORF7b as the necessary codons are not present in ORF7b gene. Among the six codons, the codon TCA has been chosen to code Serine(S) in ORF7b. So we observed a clear codon bias. Overall, there is strong choice of codon bias is seen as 59% of the codons are absent in the first positive frame of the ORF7b.

##### Double nucleotide usages

The Table 26 follows that the only double nucleotides CG and GG are absent from the gene sequence of ORF7b across the two genomes.

**Table 26:**
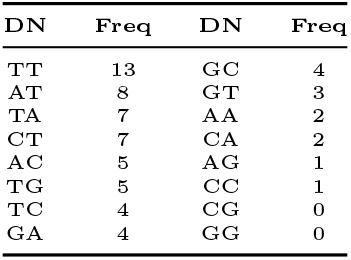
Freqeuncy of usages of double nucleotides over the gene ORF7b across the two genomes

| DN | Freq | DN | Freq |
| --- | --- | --- | --- |
| TT | 13 | GC | 4 |
| AT | 8 | GT | 3 |
| TA | 7 | AA | 2 |
| CT | 7 | CA | 2 |
| AC | 5 | AG | 1 |
| TG | 5 | CC | 1 |
| TC | 4 | CG | 0 |
| GA | 4 | GG | 0 |

The highest frequency (13) is obtained for TT in the gene ORF7b whereas the lowest frequency (1) are attained by the two double nucleotides AG and CC.

##### HEs and SEs of the spatial representations

- The spatial representations are all positively trending as the HEs are found to be greater than 0.5.
- The SEs of the spatial representations of nucleotides A, C, T and G in ORF7b are found to be same in the two genomes and they are 0.78637, 0.68404, 0.99403 and 0.55410 respectively. The SE of the binary representation of the purine-pyrimidine bases are also same and that is 0.94566 which is significantly less as compared to other genes as observed.
- The nucleotide conservation SE of the gene ORF7b across the two genomes NC_045512 and MT358637 are found to be non-identical and they are 0.91796 and 0.98152 respectively. The uncertainty of nucleotide conservation over the gene ORF7b in the genome MT358637 is higher which implies the nucleotides in ORF7b of the genome NC_045512 is conserved more than that of the other.

### 2.2. Phylogenetic Relationships of the Genomes

Based on the feature vectors obtained for each gene over the four genomes NC_045512, MT012098, MT050493 and MT358637, pairwise Euclidean distances have been enumerated. Based on the distance matrix with respect to the each gene of the four genomes, a corresponding phylogeny is obtained.

There are six different phylogenetic trees are derived from the features of respective genes of the four genomes. The phylogeny based on the features of the gene E, ORF3a, ORF6 and ORF7a are identical as shown in the Fig.8. The Fig.8 shows two phylogenetic trees-one for four genes E, ORF3a, ORF6 and ORF7a and other for two genes M and N. Interestingly, phylogenetic tree for ORF1, ORF8 and ORF10 and S are different from each other (Fig.9).

**Figure 8:**
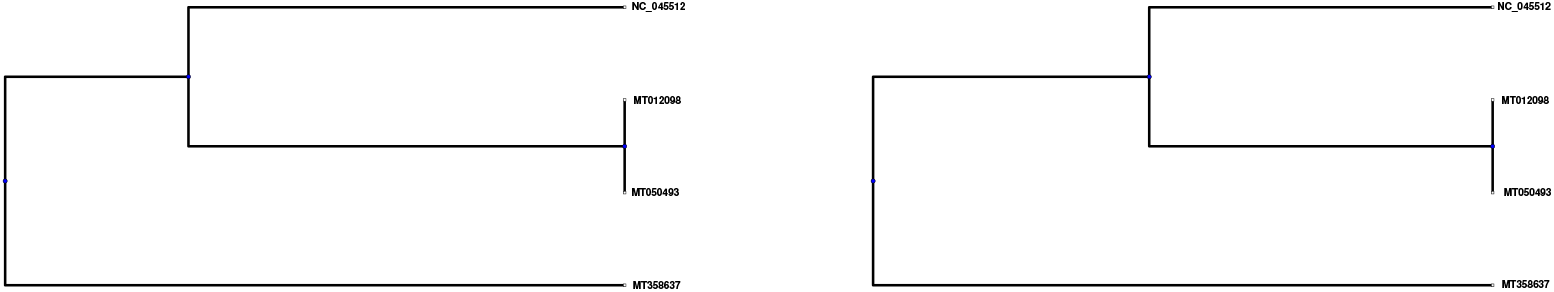
Phylogeny tree among based on the features of the genes E, ORF3a, ORF6 and ORF7a (left) and the genes M, N (right) over the four genomes.

**Figure 9:**
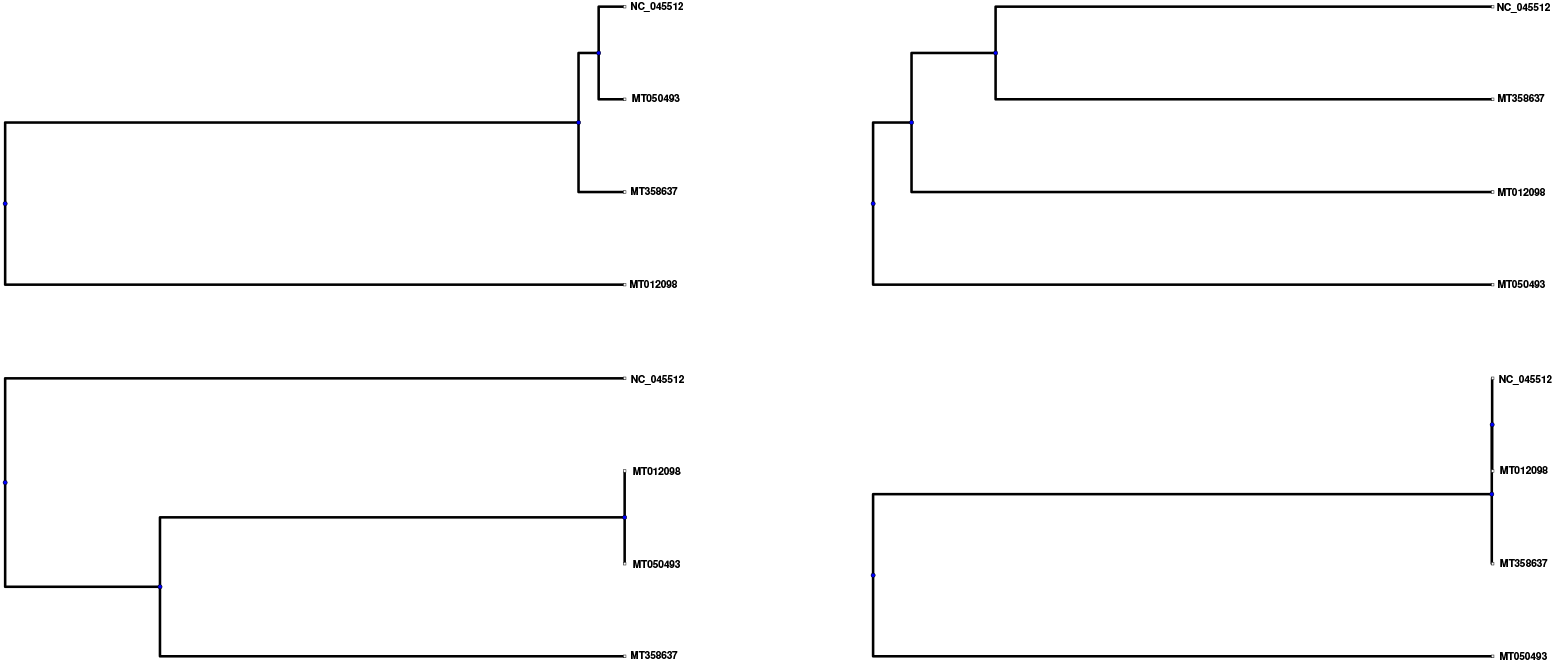
Phylogeny trees based on the features of the gene S (up-left), ORF1(up-right), ORF10 (down-left) and ORF8 (down-right) over the four genomes.

The phylogenetic relationship (Fig.3) among the four genomes, it is derived that the genomes NC_045512, MT012098, MT050493 are close enough with respect to spatial and molecular organizations of the genes M and N as compared to the other genome MT358637 which belong to the other branch of the binary tree. Here again, the sequence-similarity based phylogeny (Fig.2) is not linearly reflected while phylogeny is derived by accounting the spatial and molecular organizations of the genes M and N.

The distinct phylogenetic relationships are developed in the Fig.9 by the features of the genes ORF1, ORF10, ORF8 and S.

From the phylogeny based on the features of the gene S as shown in Fig.9, it is found that the genomes NC_045512 and MT050493 are most close to each other as belong to same level of the phylogeny. These two genomes are close to the genome MT358637. The genome MT012098 is distantly close to the other genomes since this genomes belong to a branch of primary binary level of the phylogeny. This phylogeny discriminates the genome MT012098 from NC_045512 according to the spatial and molecular organization of the gene S, although they are sequentially very close to each other as mentioned previously. The gene ORF1 is discriminating the genome MT050493 from others as depicted in the phylogeny based on the features of the gene ORF1. Further it discloses the closeness among the three other genomes NC_045512, MT012098 and MT358637. The genomes NC_045512 and MT358637 are the closest as shown in Fig.9 (up-right). In the other branch of same level of the phylogenetic tree, the genome MT012098 belongs.

As per the phylogenetic relationship based on the features of ORF10, it is found that the genome MT012098 and MT050493 are most close to each other as they below to a binary branches in the same level. Then the genome MT358637 is closed in the upper level of binary tree of the phylogeny. The genome NC_045512 is distantly related to the cluster of three other genomes. It is worth noting that the SEs of conservation of nucleotides in the gene ORF10 are the determining features of closeness among the genomes.

From the phylogenetic relationship among the four genomes based on the features extracted for the gene ORF8 it is found that the three genomes NC_045512, MT012098 and MT358637 are close enough each other as they all belong to a single branch of the binary phylogenetic tree whereas the genome MT050493 belongs to the other branch. It is observed that the Spatial arrangements of the nucleotide bases C and T in the gene ORF8 make the genome MT050493 different from other three.

It is observed from the phylogeny in the Fig.2, the genomes MT012098, MT050493 are most close to each other with respect to the features obtained for the genes E, ORF3a, ORF6 and ORF7a. These two genomes are close then with NC_045512 since they belong under the same binary branches. These three genomes are distantly close to MT358637. It is worth noting that the phylogeny based on sequential similarities among the four genomes does not go simply with the phylogeny based on the spatial features.

Differences in phylogenetic tree arrangement with these genes suggest that three genome of Indian have come from three different origin or evolution of viral genome is very fast process. Irrespective of the evolution, subtle biasness towards the usage of codon remains.

## 3. Conclusions

- There are several orders of nucleotide frequencies in different gene sequences have been observed. The pyrimidine-rich sequences E, M, S, ORF3a, ORF7a, ORF7b and ORF10 possess the order T-A-C-G. An exception happens in the case of purine-rich sequence ORF1. i.e. the order becomes T-A-C-G. The gene ORF8 being pyrimidine-rich is having the changed order as T-A-G-C. Although the N gene is purine-rich here we find a different ordering A-C-G-T while the purine-rich sequence ORF6 has the order A-T-C-G. Among all the eleven genes embedded in the four genomes the highest pyrimidine-rich (63.64%) sequence is of the gene ORF7b.
- The GC content for all these genes are widely varied over the closed interval [27.95, 47.20]. Note that both the genes, ORF6 and N having lowest and highest GC content respectively, present over all the SARS-CoV2 genomes. One may note that the GC content (47.2%) of N is significantly high which may distinctly characterize it from the other structural proteins E (38.15%), M (42.6%) and S(37.3%).
- The distribution of purine and pyrimidine bases over each gene across four genomes are found to be highly uncertain. That is, the purine and pyrimidine bases are equally likely to appear in the sequences. Although it is noted that these purine-pyrimidine spatial organizations is positively trending.
- In most of the genes all the sixteen double nucleotides are present. That is, no choice bias of double nucleotides is emerged in all the gene across the four genomes. The genes E, ORF10, ORF6 do not contain the double nucleotides GG, CC and CG respectively. In exception, the gene ORF7b does not contain two of the double nucleotides viz. CG and GG. It is noted that the gene ORF7b does not belong to the genomes of S-type as it seems from the present data. The gene ORF7b (length:132 bases) is uniquely characterized by the absence of two double nucleotides CG and GG as observed. On the other hand the ORF10 which is of the smallest length (117 bases) is characterized by the absence of only one double nucleotide CC.
- From the Table 3, it is to be pointed out that the higher consistently nucleotide conservations (SE:0.98152) over the all genes is observed in the genome MT358637 (India-Gujrat).
- Though the three Indian genomes show 99% identity with the China-Wuhan genome, phylogenetic trees are different with respective to individual genes. This suggests that recombination has more impact than mutation in evolving SARS-CoV2 on a geocentric basis in India.

Uncertainty on the distribution of purine and pyrimidine bases over each gene across the four genomes further suggest that possibility of recombination among coronavirus genus exists for evolution of SARS-CoV2 in India which may have different virulence behaviour.

## Author Contributions

SH and PPC conceived the problem. AM and RKR coded and produced the results. SH, PPC, SSJ, PP analysed the data and PPC, PP and SSJ supervised the project. SH wrote the initial draft which was checked and edited by all other authors to generate the final version.

## Acknowledgement

Authors acknowledge the financial support from IACS.

